# Immune modulatory effects of probiotic *Streptococcus thermophilus* on human monocytes

**DOI:** 10.1101/2020.08.27.271346

**Authors:** Narges Dargahi, Joshua Johnson, Vasso Apostolopoulos

**Affiliations:** Institute for Health and Sport, Victoria University, Melbourne, VIC, Australia; Institute for Sustainable Industries and Liveable Cities, Victoria University, Melbourne VIC, Australia

**Keywords:** Probiotics, microbiome, Lactic acid bacteria, *Streptococcus thermophilus*, Peripheral blood mononuclear cells, Monocyte, RNA, Innate immune response, Adaptive immune response, Inflammation

## Abstract

Ingesting probiotics contributes to the development of a healthy microflora in the gastrointestinal tract with established benefits to human health. Some of these beneficial effects may be through modulating of the immune system and probiotics have become more common in the treatment of many inflammatory and immune disorders. We demonstrate a range of immune modulating effects of *Streptococcus thermophilus* by human monocytes, including, decreased mRNA expression of IL-1R, IL-18, IFNγR1, IFNαR1, CCL2, CCR5, TLR-1, TLR-2, TLR-4, TLR-5, TLR-6, TLR-8, CD14, CD86, CD4, ITGAM, LYZ, TYK2, IFNR1, IRAK-1, NOD2, MYD88, ITGAM, SLC11A1, and, increased expression of IL-1α, IL-1β, IL-2, IL-6, IL-8, IL-23, IFNγ, TNFα, CSF-2. Routine administration of *Streptococcus thermophilus* in fermented dairy products, and their consumption may be beneficial to the treatment/management of inflammatory and autoimmune diseases.

## 1. Introduction

The human body and, in particular, the gastrointestinal tract (GIT) hosts a variety of microbial populations collectively referred to as the microbiome [1]. The GIT microbiome plays a fundamental role in the maintenance of a healthy immune system [1, 2], and any disruption to the microbiome can lead to serious ill health effects [3, 4]. In order to maintain a healthy microbiome, regular ingestion of probiotic supplements either as capsules or in fermented dairy products has been suggested. These practices have led to various improved health outcomes and treatment of ill health, such as infections, constipation and diarrhoea [1, 5, 6].

The majority of probiotics belong to the lactic acid bacteria (LAB) family; gram positive lactic acid producing microorganisms that include several genera such as bifidobacteria, lactobacilli streptococci and enterococci [1]. The small and large intestines are highly populated with these microorganisms [7-9], and are routinely supplemented in foods as live strains due to their established beneficial effects to human health [1, 2, 9-14]. *Streptococcus* species such as exopolysaccharide-producing strains of *Streptococcus thermophilus* (ST) [13, 15, 16] are amongst those consumed. ST is used for fermentation of milk products and is recognized as an important species for its health benefits [17, 18]. In fact, ST and *L. brevis* synergistically display health benefits which are well established, also, ST is one of the bacteria in the VSL#3 probiotic mixture, which has been applied for the treatment of inflammatory conditions [19, 20]. Probiotics also interact with the immune system where they exhibit immunomodulatory and anti-inflammatory effects [3, 21, 22].

Use of probiotic bacteria can increase the abundance of and concurrently modulate immune cells including B, T helper (Th)-1, Th-2, Th-17 and regulatory T (Treg) cells. This in turn, directly influences human health and modulates pathologies of immune/autoimmune diseases [1, 2, 14]. In fact, primary macrophages co-cultured with ST bacteria have been shown to increase production of anti-inflammatory IL-10 and pro-inflammatory IL-12 cytokines [23]. ST1275 and *Bifidobacterium longum* BL536 induce expression of high levels of transforming growth factor (TGF)-beta, a key factor in the differentiation of Treg and Th-17 cells by bulk peripheral blood mononuclear cell (PBMC) cultures [24]. Probiotic bacteria, however, can only confer these benefits through interaction with specific immune cells, primarily antigen presenting cells (APC), which include monocytes, as mediators between bacteria/foreign agents and the immune system’s effector adaptive immune cells [25].

In line with these findings, we previously noted that ST1342, ST1275 and ST285 modulated U937 monocyte cell line by increasing IL-4, IL-10, GM-CSF and CXCL8 production. In addition the cell surface marker expression of CD11c, CD86, C206, CD209, MHC-1 were upregulated, suggesting that ST bateria has an influence on the immune system [1]. Furthermore, we recently showed that ST285 exerted an array of anti-inflammatory immune-modulatory properties to human PBMC [26]. In particular, ST285 decreased mRNA expression of IL-18, IFNγR1, CCR5, CXCL10, TLR-1, TLR-2, TLR-4, TLR-8, CD14, CD40, CD86, C3, GATA3, ITGAM, IRF7, NLP3, LYZ, TYK2, IFNR1, and upregulated IL-1α, IL-1β, IL-6, IL-8, IL-10, IL-23, IFNγ, TNFα, CSF-2 [26]. The data demonstrated a predominant anti-inflammatory profile exhibited by ST285. Due to the role of monocytes and their progeny in initiation and maintenance of both innate and adaptive immune responses, we now show immune modulatory properties of ST285 on monocytes from healthy blood donors. The data paves the way for further work to determine the effects of ST285 in inflammatory disease models *in vitro* and *in vivo*, such as multiple sclerosis, inflammatory bowel disease and allergies.

## 2. Material and methods

### 2.1. Bacterial strains

Pure bacterial cultures of ST285 were obtained from Victoria University culture collection (Werribee, VIC, Australia). Stock cultures were stored in cryobeads at −80° C. Prior to each experiment the cultures were propagated in M17 broth (Oxoid, Denmark) with 20 g/L lactose and incubated at 37° C under aerobic conditions. In order to confirm gram-positivity and assess purity, morphology and characteristics, bacteria were cultured in M17 agar (1.5 % w/v agar) with 20 g/L lactose (Oxoid, Denmark) as well [1].

### 2.2 Preparation of live bacterial suspensions

Prior to experiments bacteria medium was prepared and autoclaved at 121° C for 15 minutes (mins) and bacterial cultures were grown 3 times in M17 broth with 20 g/L lactose, at 37° C aerobically for 18 hours (hr) with a 1 % inoculum transfer rate [27] at 37-42° C [15]. Bacteria were harvested during stationary growth phase on the day of experiment, centrifuged (6000×g) for 15 min at 4° C, followed by washing twice with phosphate-buffered saline (PBS) (Invitrogen, Pty Ltd. Australia) and resuspended in the Roswell Park Memorial Institute (RPMI) 1640 culture media (Invitrogen, Pty Ltd. Australia), which constituted the live-bacteria suspensions.

### 2.4. Enumeration of bacterial cells

Prior to co-culturing with PBMC, bacterial strains cultured in M17 broth, were centrifuged and transferred into PBS (Invitrogen, Pty Ltd. Australia), adjusted to a final concentration of 10^8^ colony forming units (cfu)/ml by measuring the optical density at 600 nm. Then washed twice with PBS and resuspended in RPMI 1640 (Invitrogen, Pty Ltd. Australia) [1].

### 2.4. Isolation of monocytes from buffy coat

Buffy coats were received from the Australian Red Cross blood bank in Melbourne, and PBMC were isolated using standard Ficoll-Paque density gradient centrifugation method as previously described [11]. PBMC cells were resuspended at ∼5 × 10^8^ cells/mL in adequate amount of Dulbecco’s phosphate-buffered saline, D-PBS (D-PBS without Ca^++^ and Mg^++^) supplemented with 2% FBS and 3 mM cell culture grade EDTA (Life Technologies; Thermofisher) prior to monocyte isolation. Monocytes were isolated using the EasySep Human Mono Isolation Kit (STEMCELL technology, Canada) [28]. Isolation method involved the use of immunomagnetic negative selection method targeting CD16^+^ monocytes, excluding non-monocyte cells, and platelets, yielding highly pure CD14^+^CD16^-^ monocytes. As such the unwanted cell populations are labelled with specific cell surface marker antibodies and magnetic particles, and removed following separation by using an EasySep™ magnet (STEMCELL technology, Canada) according to manufacturer’s instructions [28]. Monocyte cells were added into a fresh tube, checked for viability and purity.

### 2.5. Stimulation of monocytes with ST285

Monocytes (∼3-5× 10^7^ cells) isolated from three different donors were resuspended in RPMI 1640 media supplemented with 10% heat-inactivated FBS (Invitrogen, Pty Ltd. Australia), 1% antibiotic-antimycotic solution and 2 mM L-glutamine in cell culture flasks, into which 5×10^8^ ST285 bacteria were added. Monocytes (∼3-5× 10^7^ cells) minus the ST285 bacteria were used as a control and incubated at 37° C, 5 % CO_2_ for 24 hrs [1]. In previous studies we demonstrated that 24 hrs co-culture was optimal for stimulation of the U937 monocyte cell line, and all incubations described herein were for 24 hrs [1]. Monocytes were harvested post incubation period, snap frozen and stored at −80° C.

### 2.6. RNA extraction from monocytes

Total RNA was extracted from stimulated and unstimulated monocytes using the RNeasy® mini kit (Qiagen, Hilden, Germany) according to the manufacturer’s instructions. Briefly, monocytes were harvested using centrifugation, supernatants were removed and RNA was extracted from each pellet by resuspending pellet in lysis buffer supplemented with β-mercaptoethanol for cell disruption. Monocytes were lysed and each cell lysate was homogenized by passing through Qia-shredder columns (Qiagen, Hilden, Germany). Each monocytes lysate was then mixed 1:1 with 70% ethanol (equal volume) and were transferred onto RNeasy mini-spin columns. DNA was eliminated using DNase digestion/ treatment using RNase-Free DNase Set (Qiagen, Hilden, Germany) by adding it directly onto the columns. The RNA Integrity Number (RIN) of all RNA samples were determined using an Agilent 2100 Bioanalyzer and Agilent RNA 6000 nano kit (Agilent Technologies, Santa Clara, CA, USA). A minimum RIN of 7.5 was used as the standard for inclusion in the gene expression study. Subsequently, the concentration of each individual monocyte RNA sample was quantified using a Qubit RNA BR Assay (Invitrogen, Pty Ltd. Australia).

### 2.7. Assessing changes in the expression of genes associated with innate and adaptive immunity

Using RT^2^ first strand kit (Qiagen, Hilden, Germany), adequate aliquots of each RNA sample was reverse-transcribed to produce complementary DNA (cDNA) according to the manufacturer’s instructions. Quantitative real-time polymerase chain reaction (qRT-PCR) was carried out by using the ‘Human Innate and Adaptive immune Response’ kit (Qiagen, Hilden, Germany) to assess the expression of genes/mRNA. Using a CFX Real-Time touch PCR System thermo-cycler (Biorad, Melbourne Australia) and Qiagen prescribed cycle, the relative gene/mRNA expression of ST285-treated monocytes were analyzed in contrast to control untreated monocytes. The RT^2^ qPCR innate and adaptive immune response arrays targeted a set of 84 innate and adaptive immune-related genes, five housekeeping genes, an RT control, a positive PCR control, and a human genomic DNA contamination control [29]. Relative gene expression was calculated using the Qiagen webportal PCR array data analysis web-based software (Qiagen, Germany). Differential expression (up and down regulation) of the genes were identified using the criteria of a > 2.0-fold increase/decrease in gene expressions in treated monocytes in comparison with those genes in control monocyte cultures.

### 2.8. Data analysis

The Delta-Delta CT (ΔΔCT) method was used for calculating fold-changes [30]. Fold-regulation represents fold-change results in a biologically meaningful way. In these RT2 profiler PCR array results, fold-change values >1, indicate a positive (or an up-) regulation. Actually, in the case of genes which are upregulated, the fold-regulation is equivalent to the fold-change. Fold-change rates <1 indicate a negative (or a down) regulation. In the case of negative values, the fold-regulation is actually the negative inverse of the fold-change [31-33]. Data related to changes in the expression of the genes were estimated using Qiagen RT^2^ profiler data analysis webportal that uses the ΔΔCT method in calculating fold-changes. The raw CT values were uploaded to the Qiagen data analysis webportal with the lower limit of detection set for 35 cycles and 3 internal controls. For controls, RT efficiency, PCR array reproducibility, and genomic DNA contamination were assessed to ensure all arrays successfully passed all the control check-ponits. Normalization of the raw data was done by using the incorporated housekeeping genes (HKG) panel. Then using the ΔΔCT method, both housekeeping gene references and controls (untreated monocytes in RPMI) were evaluated to determine relative expression of mRNA.

### 2.9. Statistical analysis

The p values were calculated by the use of a Student’s *t-test* of the Triplicate 2^ (-Delta CT) [(2^-ΔCT)] values for each gene in treatment groups (monocyte co-cultured with ST) and the control group (monocyte in RPMI media) [31, 32].

## 3. Results

Among 84 genes evaluated, expression of 30 genes were significantly altered with over 2.0 fold up or down regulations in monocyte samples (n = 3) following co-culture with ST285 compared to control (Figure 1).

**Fig. 1.**
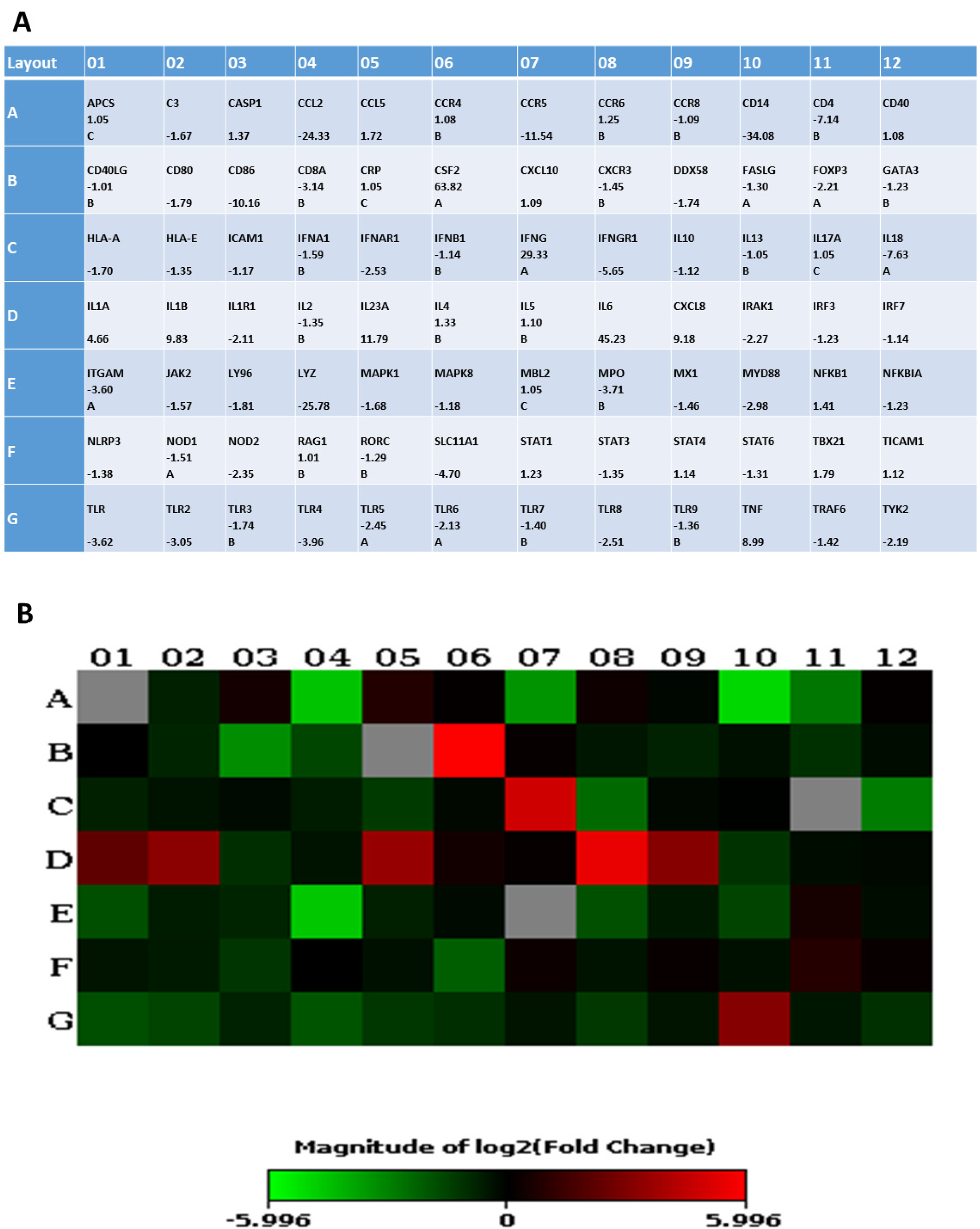
Effects of co-culturing ST285 with monocytes (n=3) on gene/RNA expression compared to control monocytes after 24 hrs. (A) All 84 genes are shown including those with significant high up/down regulated genes (more than 2-fold) and those with no significant change (less than 2-fold). The housekeeping genes (HKG) panel and other genes used for normalization of the raw data are not presented. In case of no letter or comments, **t**he expression of gene/s is relatively high in both the test and control group (threshold cycle (CT) is <30). Letter A specifies the gene’s average threshold cycle to be reasonably high (> 30) in either the treated samples or the controls and relatively low (< 30) in the other/opposite sample. Thus, in case of presenting fold changes with letter A, the estimate fold change may be an underestimate. Letter B suggests a reasonably high (> 30) gene’s average threshold cycle that means a low level of average expression of relevant gene, in both test/treated samples and untreated control samples, and the p-value for the fold-change might be either relatively high (p > 0.05). Thus, in case of presenting fold changes with letter B, the estimate fold change may be slightly overestimate or unavailable. Letter C indicates that that gene’s average threshold cycle is either not determined or greater than the defined default 35 cut-off value, in both test/treated samples and control samples, suggesting that its expression was not detectable, resulting in the fold-change values being un-interpretable [72-74]. (B) Presentation of data as a heatmap of average gene/RNA expressions of monocytes (n=3) co-cultured with ST285, compared to control. Green represents down regulated genes to red represents upregulated genes.

### 3.1. ST285 alters cytokine gene expression levels of monocytes

#### 3.1.1. ST285 causes upregulation of IL-1α, IL-6 and IL-23 and downregulation of IL-1R1 genes

IL-1α was upregulated 4.66 ± 0.7 fold, IL-1β was upregulated 9.83 ± 0.49 fold, IL-6 was upregulated 42.23 ± 0.32 fold and IL-23α was upregulated 3.8 ± 1.0 fold (Figure 2). IL-1R1 was downregulated 2.11 ± 0.36 fold (Figure 2). Neither IL-17A nor IL-2, and IL-10 were altered following monocyte co-cultured with ST285.

**Fig. 2.**
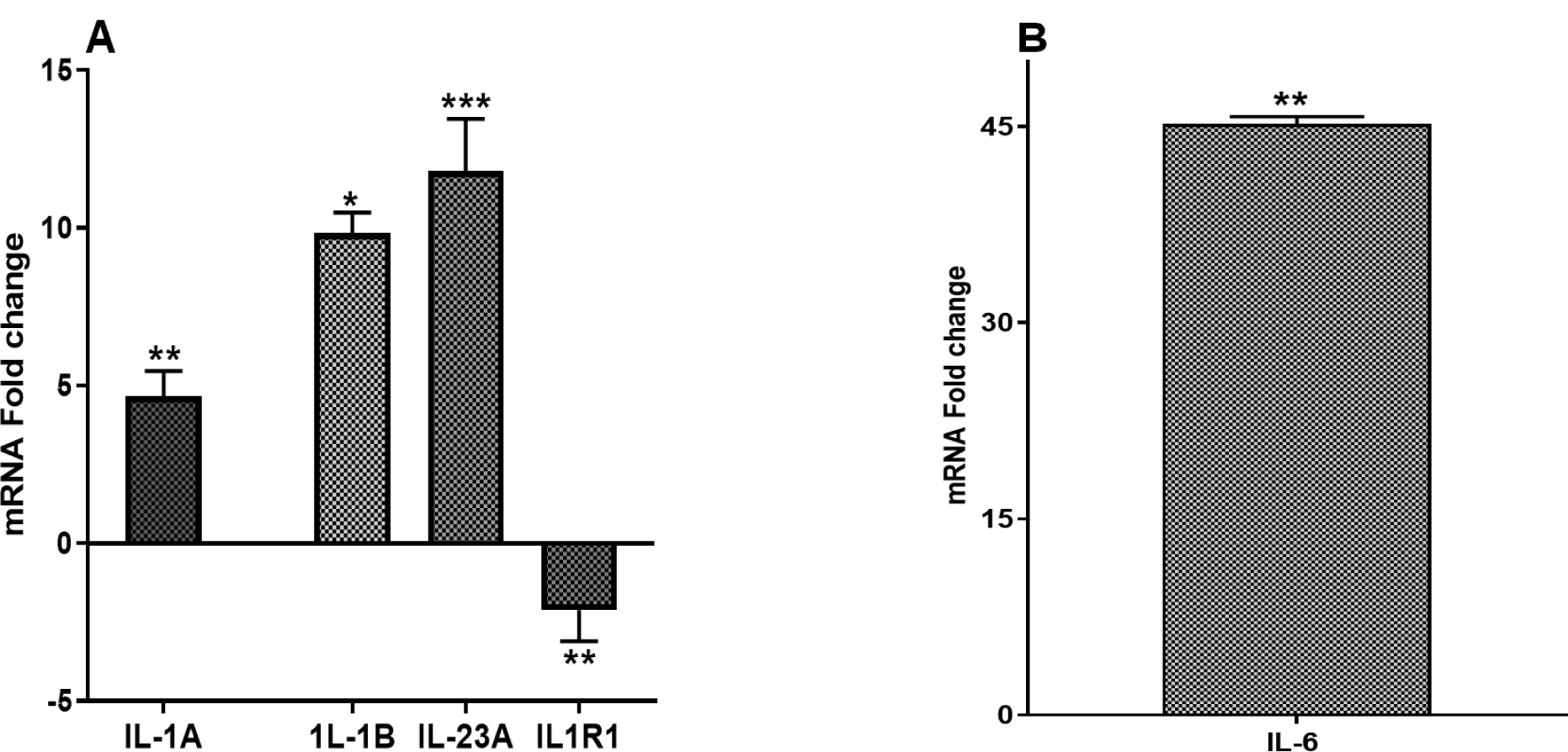
(A) IL-1α, IL-1β, IL-23α, IL-1R1 and (B) IL-6, mRNA fold change following 24 h co-culture of ST285 with monocytes (n=3), compared to control monocytes. The innate and adaptive RT^2^ gene profiler arrays were used to determine changes in gene expression. Symbols represent *p* value for Tukey Test (One way ANOVA) where ** *p* < 0.04 and *** *p* < 0.02.

#### 3.1.2. Modulation of pro-inflammatory cytokines

ST285 induced upregulation of IFNγ (29.33 ± 0.26 fold) (Figure 3A). IL-18 a Th1 inducing pro-inflammatory cytokine was downregulated (7.63 ± 0.37 fold) (Figure 3A). In addition, IFNγR1, a transmembrane protein which interacts with IFNγ, was also downregulated 5.65 ± 0.05 fold and IFNAR1 (involved in defence against viruses) was downregulated 2.53 ± 0.05 fold (Figure 3A). Tumor-necrosis factor-alpha (TNFα), which is important in the defense against bacterial infections, and in acute phase reactions was upregulated 8.99 ± 1.06 fold (Figure 3). Gene expressions of other cytokines, IFNA1, IFNB1, IL-4, IL-5, IL-12 and IL-13 were not significantly altered.

**Fig. 3.**
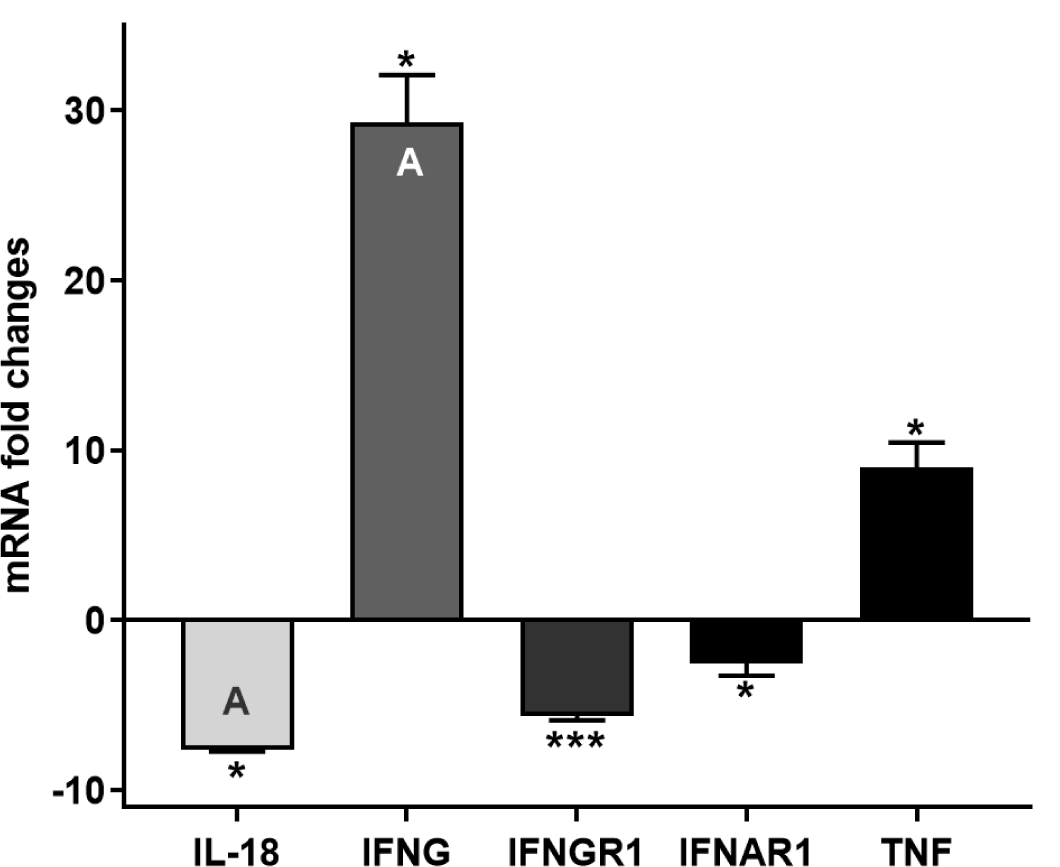
IL-18, IFN-γ, and IFN-γ R1, IFN-α R1 and TNF mRNA mRNA fold change following 24 h co-culture of ST285 with monocytes (n=3), compared to control monocytes. The innate and adaptive RT^2^ gene profiler arrays were used to determine changes in gene expression. Symbols represent *p* value for Tukey Test (One way ANOVA) where * *p* < 0.05 and *** *p* < 0.02.

### 3.2. ST285 alters chemokine gene expression levels of monocytes

CCR5 a Th1 marker involved in immune response and CCL2 (MCP-1) involved in humoral immunity were down regulated 11.54 ± 0.23 and 24.33 ± 1.44 fold respectively (Figure 4). Chemokine (CXCL8, IL-8), important in the innate immune system, stimulates chemotaxis, was upregulated 9.18 ± 0.26 fold following ST285 co-culture with monocyte cells (Figure 5). However, no significant differences were noted for gene expressions of other chemokines, including CXCL10 (INP10), CCL5 (RANTES), CCL8, CCR4, CCR8 and CXCR3 following monocytes’ exposure to ST285.

**Fig. 4.**
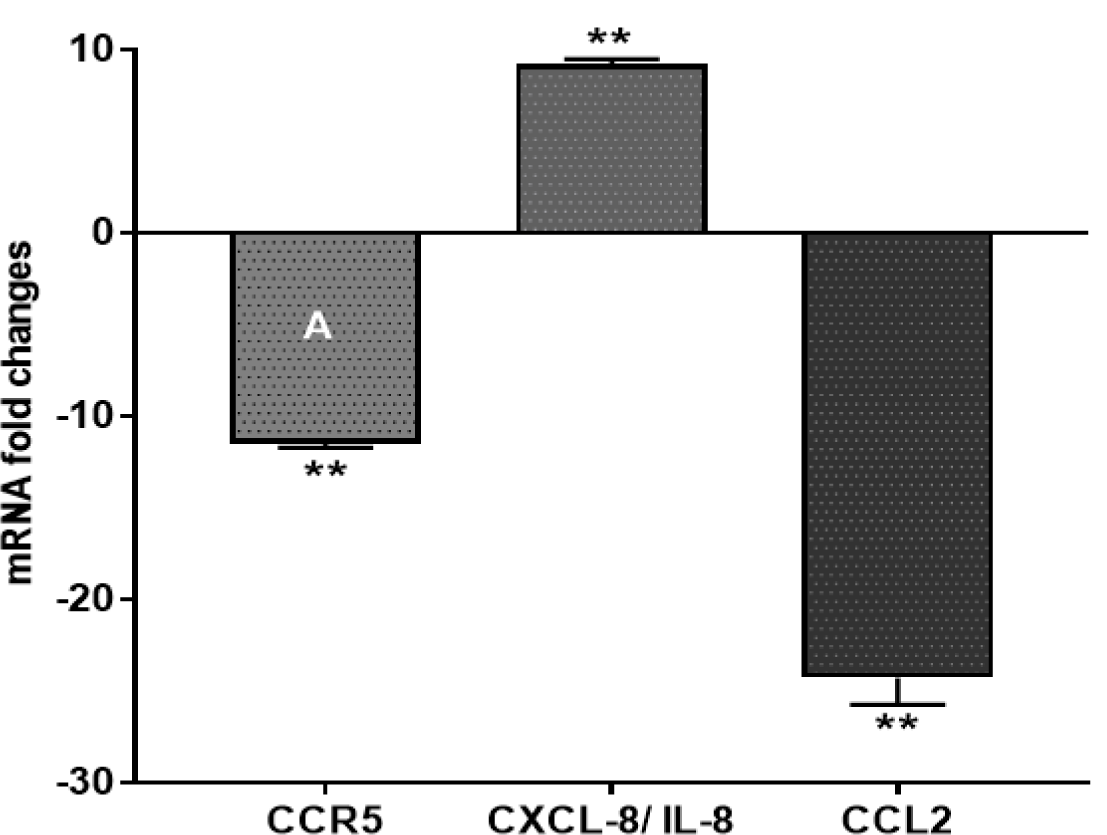
CCR5, CXCL8 (IL-8) and CCL2 mRNA fold change following 24 h co-culture of ST285 with monocytes (n=3), compared to control monocytes. The innate and adaptive RT^2^ gene profiler arrays were used to determine changes in gene expression. Symbol represents *p* value for Tukey Test (One way ANOVA) where ** *p* < 0.04.

**Fig. 5.**
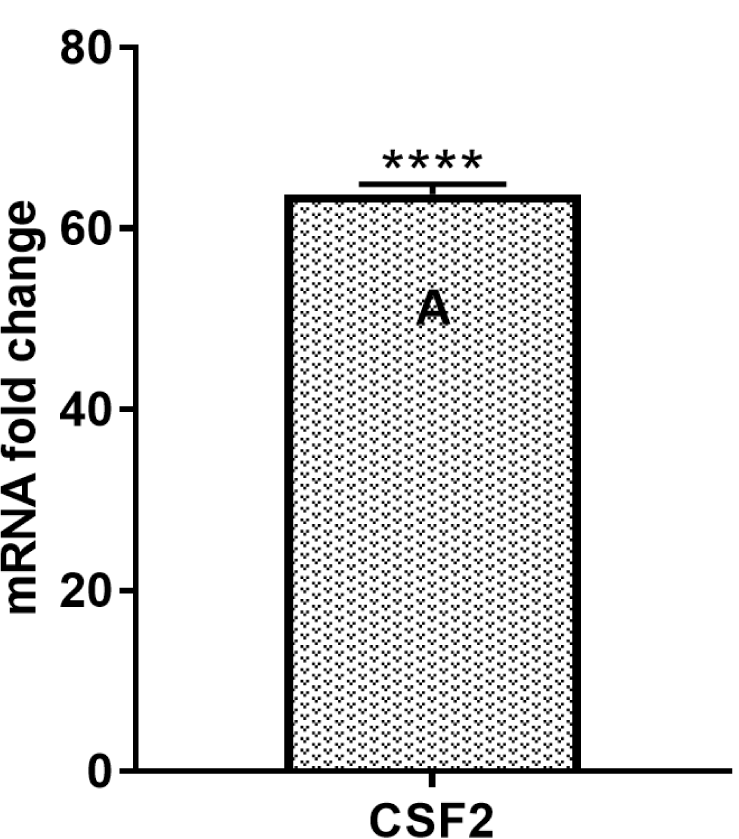
CSF-2, mRNA fold change following 24 h co-culture of ST285 with monocytes (n=3), compared to control monocytes. The innate and adaptive RT^2^ gene profiler arrays were used to determine changes in gene expression. Symbol represents *p* value for Tukey Test (One way ANOVA) where **** p < 0.01.

### 3.3. Significant upregulation of colony stimulating factor mRNA expression levels

Colony-stimulating factor (CSF)-2 which enables cell proliferation and differentiation of cells, was significantly increased by 63.82 ± 1.12 fold (Figure 5) after co-culturing monocytes with ST285 bacteria.

### 3.4. ST285 alters Toll like receptor gene expression levels of monocytes

TLR (toll like receptor)-1, TLR-2, TLR-4, TLR-5, TLR-6 and TLR-8 are part of the innate immune response and involved in the defense response to bacteria. Monocytes co-cultured with ST285 induced significant differential downregulation of TLRs; TLR-1 (−3.63 ± 0.14), TLR-2 (−3.05 ± 0.36 fold), TLR-4 (−3.96 ± 0.16 fold), TLR-5 (−2.45 ± 0.23 fold), TLR-6 (−2.13 ± 0.23 fold), and TLR-8 (−2.51 ± 0.12 fold) (Figure 6). However, changes to TLR-3 and TLR-9 were not significant.

**Fig. 6.**
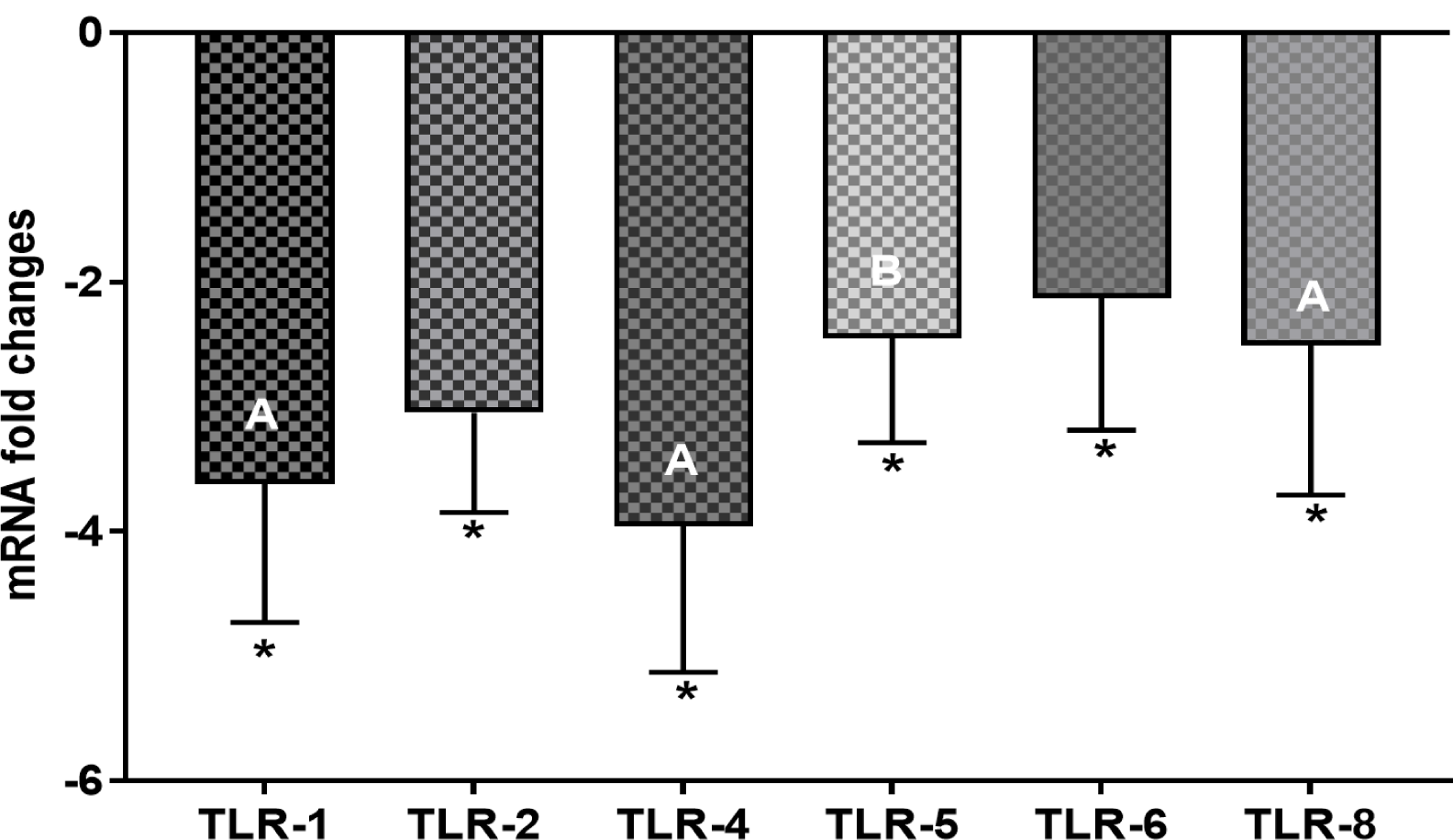
TLR-1, TLR-2, TLR-4, TLR-5, TLR-6 and TLR-8, mRNA fold change following 24 h co-culture of ST285 with monocytes (n=3), compared to control monocytes. The innate and adaptive RT^2^ gene profiler arrays were used to determine changes in gene expression. Symbol represents *p* value for Tukey Test (One way ANOVA) where * *p* < 0.05.

### 3.5. Cell surface markers CD14, CD86 and CD4 mRNA expression levels

Expression of the monocyte cell surface markers CD14 and CD86 were significantly downregulated 34.08 ± 3.42 and 10.16 ± 0.14 fold, respectively (Figure 7). CD4 is expressed by Th cells, monocytes, macrophages (MQ), and dendritic cells (DCs), was downregulated 7.14 ± 0.41 fold. No significant change was observed in the expression of CD8A, CD40, CD80, GATA3, FOXP3, STAT3, CD40LG (TNFSF5), HLA-A, HLA-E and RORC genes.

**Fig. 7.**
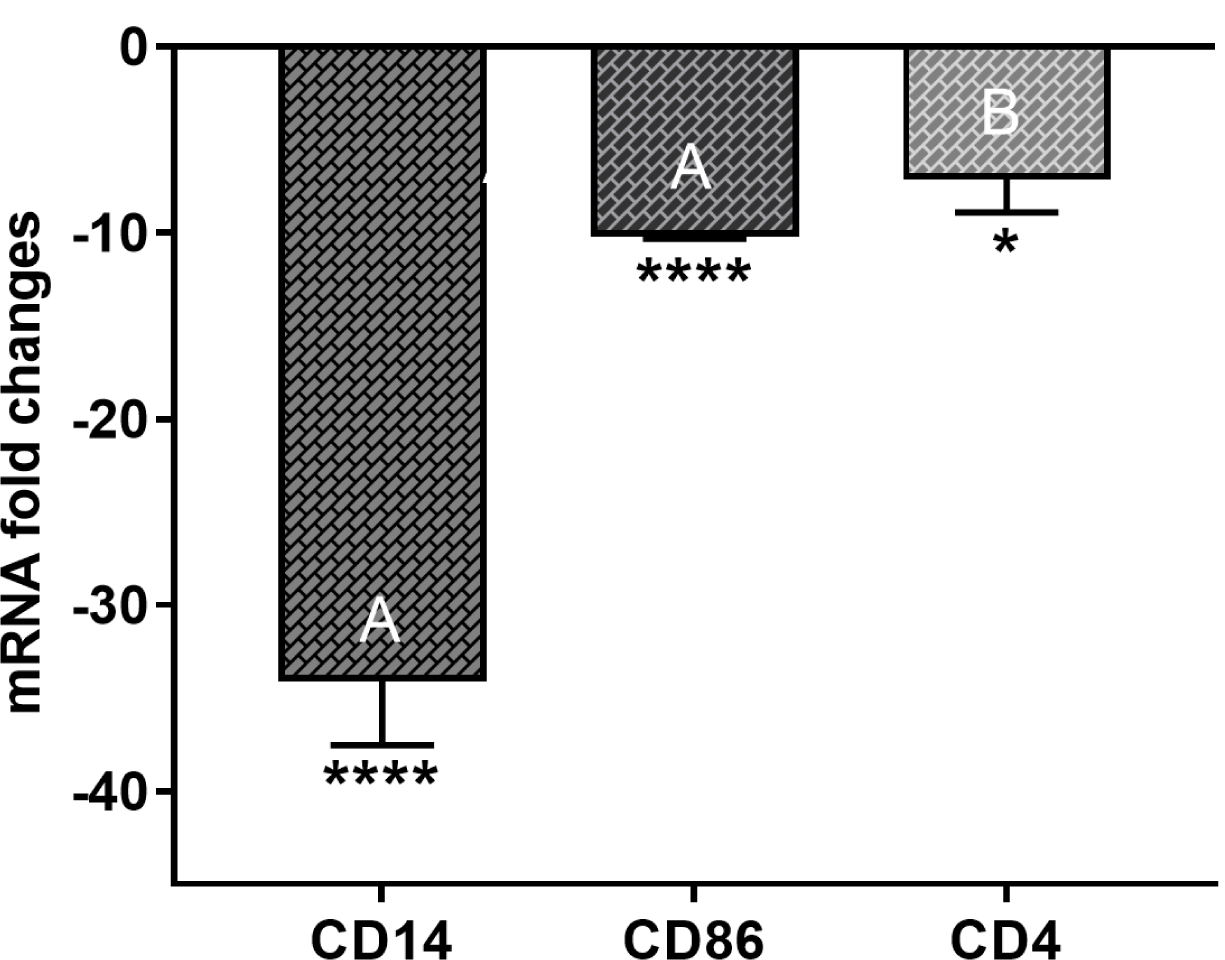
CD14, CD86 and CD4 mRNA fold change following 24 h co-culture of ST285 with monocytes (n=3), compared to control monocytes. The innate and adaptive RT^2^ gene profiler arrays were used to determine changes in gene expression. Symbols represent *p* value for Tukey Test (One way ANOVA) where * *p* < 0.05 and *** *p* < 0.02.

### 3.6. Changes to other innate and adaptive molecules, mRNA expression levels

Altered expression levels are noted in other genes following ST285 co-culture with monocytes. Significant downregulation of the following genes were noted: TYK2 (−2.19 ± 0.37), IRAK-1 (−2.27 ± 0.45), NOD2 (−2.35 ± 0.04), MYD88 (−2.98 ± 0.23), ITGAM (−3.6 ± 0.23), MPO (3.71 ± 0.12), SLC11A1 (−4.7 ± 0.17) (Figure 8A), and LYZ (25.78 ± 0.36) (Figure 8B). Other immune markers including FASLG (TNFSF6), ACTB, GATA3, complement component (C)-3, CRP, IFNAR1, JAK2, IL-1R1, MAPK8 (JNK1), IRF3, MBL2, NLRP3, NFKB1, MX1, ICAM1, MBL2, NOD1 (CARD4), DDX58 (RIG-I), RAG1, TICAM1 (TRIF) and IRF7 showed no significant mRNA gene changes in the levels of their expression.

**Fig. 8.**
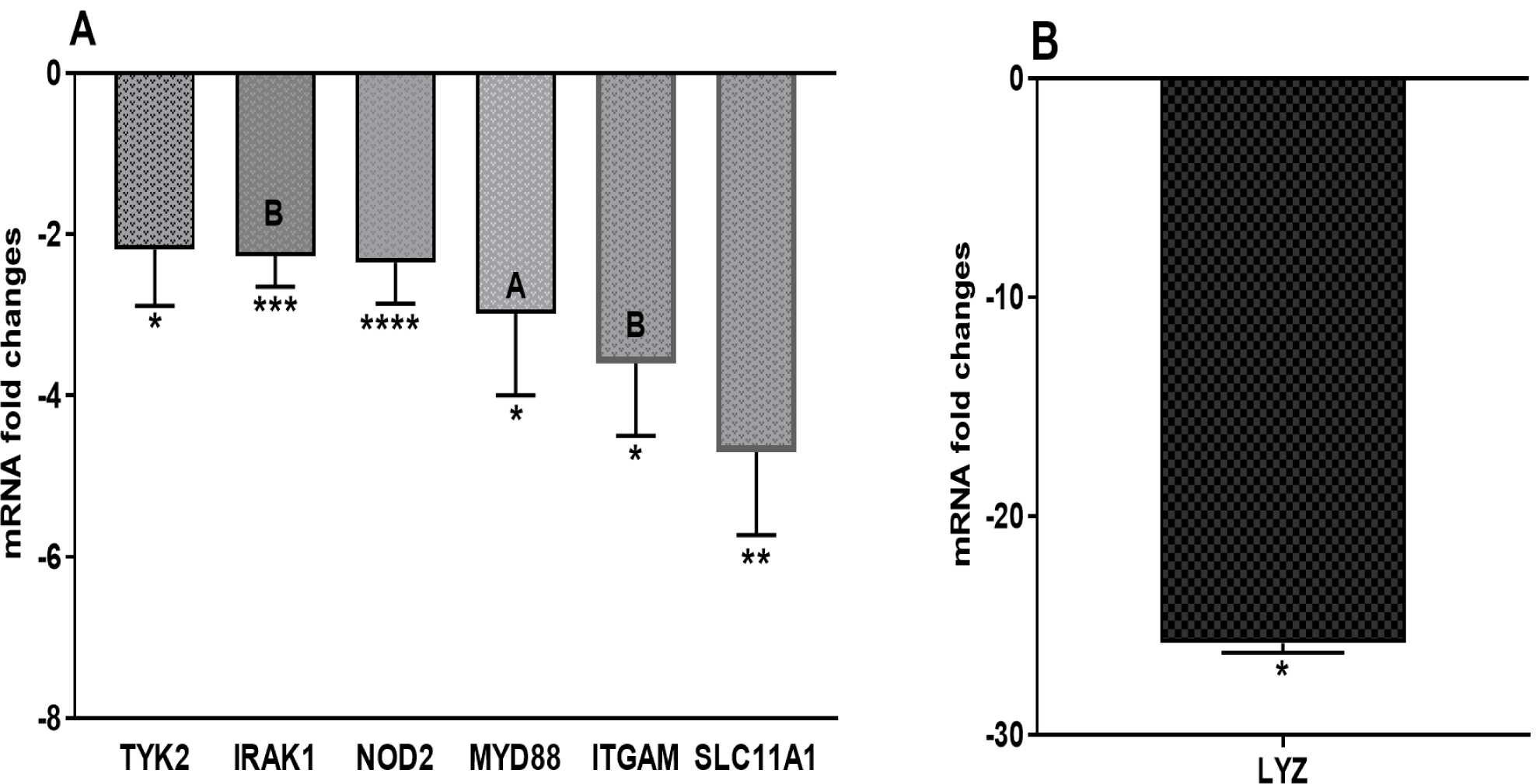
(A) TYK2, IRAK1, NOD2, MYD88, ITGAM, SLC11A1 and (B) LYZ and GATA3, mRNA fold change following 24 h co-culture of ST285 with monocytes (n=3), compared to control monocytes. The innate and adaptive RT^2^ gene profiler arrays were used to determine changes in gene expression. Symbols represent *p* value for Tukey Test (One way ANOVA) where * *p* < 0.05, ** *p* < 0.04, *** *p* < 0.02 and **** *p* < 0.01.

## 4. Discussion

ST285 co-cultured with human monocytes resulted in significant changes to 30 genes associated with different immune responses of the innate and adaptive immunity compared to control. In particular, mRNA gene expression of IL-1R, IL-18, IFNγR1, IFNαR1, CCL2, CCR5, TLR-1, TLR-2, TLR-4, TLR-5, TLR-6, TLR-8, CD14, CD86, CD4, ITGAM, LYZ, TYK2, IFNR1, IRAK-1, NOD2, MYD88, ITGAM, SLC11A1 are downregulated. Whilst ST285 increases mRNA expression of IL-1α, IL-1β, IL-1α R, IL-6, IL-8, IL-23, IFNγ, TNFα and CSF-2. These results were broadly in agreement with our previous findings showing a predominant anti-inflammatory profile by human PBMC upon co-culture with ST285 [26]. Likewise, our previous data showed a similar trend for a number of cytokine, chemokine and cell surface markers for three different ST bacteria to human U937 monocyte cell line, where ST285 was most effective [1].

### 4.1. ST285 induces IL-1α and IL-6 and downregulates IL-1R1

IL-1α secreted by DCs and MQs, usually initiates Th2 differentiation, while preventing polarization of Th1 cells [34]. IL-6 is produced by activated immune cells including monocytes/MQs [35]. IL-1α and IL-6 are significantly upregulated, whereas IL-1R1 (CD121a), a key mediator associated with several inflammatory and immune responses is downregulated in monocytes after exposure to ST285. This is in accord to PBMCs co-cultured with ST285 [26] and U937 monocyte cell line co-cultured with ST285 [1]. Likewise, it was recently noted that spleen cells from mice immunized with agonist myelin basic protein peptide (MBP_83–99_) peptide was cultured with ST285 in the presence of recall agonist peptide, which lead to significant production of IL-6 which was three times that of control without ST285 bacteria but with agonist peptide; these data suggested that ST285 has the potential to significantly change the balance towards a healthier state [36]. IL-6 acts as both pro- and anti-inflammatory cytokine [37] and its anti-inflammatory roles are associated with its inhibitory effects on IL-1, TNF-α, and activation of IL-10 and IL-1Ra [37, 38]. On the other hand, the inhibitor of NF-κB kinase (IKK) governs IL-6 mRNA stability (through phosphorylation of regnase-1), in response to IL-1R/TLR stimulation [39]. As such, *Lactobacillus paracasei* has been shown to reduce IL-6 production via prevention of NF-κB activation to THP-1 cell line [40] which is in contrast with our findings. Whereas, the surface-associated exopolysaccharide (EPS) extracted from *L. paracasei* DG showed immune-stimulating properties to human monocytic cell line THP-1 by increasing TNF-α and IL-6 gene expression which is in line with our findings [41]. In addition, human monocytes and monocyte-derived DCs co-cultured with *Veillonella parvula, Escherichia (E*.*) coli, B. adolescentis* and *L. plantarum* strains, stimulated high level of IL-6 upon exposure to *V. parvula* and *E. coli* but not *B. adolescentis* and *L. plantarum* [42].

IL-1β is secreted by monocytes and activated MQs, is involved in regulating immune and inflammatory responses to bacterial infections and injuries, hence its role in innate immunity [43]. IL-1β is upregulated by ST285 co-cultured with monocytes, which is similar to ST285 stimulation of PBMC [14], although ST285 did not stimulate IL-1β in the U937 monocyte cell line [1]. However, in other studies *L. paracasei* cultured with THP-1 cell line either before LPS treatment or together with LPS, reduced IL-1β secretion [40]. Additionally, mice immunized with agonist MBP_83–99_ peptide, spleen cells cultured with recall agonist peptide in the presence of ST285 decreased production of IL-1β [36]. Consumption of a mixed probiotic or a conventional yogurt with equal *S. thermophiles, L. bulgaricus* and surplus *L. casei* DN114001, induces high IL-1β production by *ex vivo* cultured monocytes following LPS and phytohaemmaglutinin stimulation [44].

The increased expression of IL-1α and IL-6, suggests the role of ST285 in the induction of immune responses required for acute phase (including MQs differentiation, B cell maturation, and activation of Th2 differentiation and prevention of Th1 polarization). A decrease in IL-1R1 gene expression could highlight the role of ST285 as a brake that controls the pro-inflammatory roles of both IL-6 and IL-1α.

### 4.2. ST285 changes expression of cytokines involved in inflammation and defence against bacteria

IL-18 is associated with severe inflammatory responses and plays a role in inflammatory and autoimmune disorders. Monocytes co-cultured with ST285 significantly reduced the gene expression of IL-18, which is in agreement with our recent study of ST285 co-cultured with PBMC [26], suggesting an anti-inflammatory role for ST285 bacteria. IFN-γ is an important activator of MQs, is secreted by monocytes, NK and NKT cells, and is critical for functional innate and adaptive immune responses against viruses, some bacterial and protozoa infections [45]. Monocytes co-cultured with ST285 show increased gene expression of IFN-γ suggesting an anti-bacterial response. Similarly, blood monocytes from healthy individuals who ingested either a probiotic mixed of *S. thermophiles, L. bulgaricus* and surplus *L. casei* DN114001 or a conventional yogurt containing same probiotic mixture, showed increased production of IFN-γ upon co-culturing monocyte cells *ex vivo* with LPS and phytohaemmaglutinin [44]. In another study, the effects of *L. casei* Shirota on monocyte was shown indirectly; as depletion of monocytes from PBMC co-cultured with *L. casei* Shirota was associated with an absence of IFN-γ and other cytokines demonstrating the importance of monocytes against bacterial challenge [46]. Similarly, *L. plantarum* alone and mixed *L. plantarum* and *Helicobacter pylori* added to monocytes (and lymphocytes) resulted in the production of high levels of IFN-γ with *L. plantarum* alone, compared to the mixed cultures [47]. In comparison, it was shown that IFN-γ secretion was reduced by spleen cells of mice immunized with agonist MBP_83–99_ peptide in the presence of ST285. The reduction of inflammatory IFN-γ is important in the inflamed environment situations such as inflammatory and inflammatory diseases because any level of reduction in the amount of inflammatory mediators can contribute to the relief of symptoms [36].

TNFα, a pro-inflammatory cytokine is required against bacterial infections and is involved in activating and recruiting T and B cells in the initiation of adaptive immune responses. We show upregulation of TNFα when human monocytes are co-cultured with ST285, in agreement with observations with PBMCs [26] and the U937 monocyte cell line [1]. Isolated human monocytes and monocyte-derived DCs co-cultured with *V. parvula, E. coli, B. adolescentis* and *L. plantarum* strains, similarly showed higher levels of TNFα [42]. In addition, EPS from *L. paracasei* DG also induced increased TNFα gene expression by THP-1 monocyte cell line [41]. Although, *L. paracasei* itself decreased TNF-α production by THP-1 cell line via inhibition of NF-κB activation [40]. Similarly, *L. plantarum* genomic DNA reduced the production of TNFα in THP-1 monocyte cells [48]. Additionally, the importance of monocytes in phagocytosis was shown by using monocyte-depleted-PBMC in co-culture with *L. casei* Shirota, which led to no secretion of TNFα [46]. Similarly, spleen cells from immunized with MBP_83–99_ peptide mice demonstrated marginally decreased TNFα production in the presence of ST285 and recall MBP_83–99_ peptide; marginal reduction of both TNFα and IFN-γ subtractions could be advantageous for twisting inflamed status of diseases into healthier normal status [36].

IFNAR1 is a membrane protein and a receptor for both IFNα and IFNβ associated with defence against viruses. IFNAR1 signalling is involved in production of pro-inflammatory cytokines [49], as such that IFNAR1 knockout mice demonstrate reduced pro-inflammatory chemokines and cytokines [49]. IFNAR1 is significantly downregulated by monocytes following co-culture with ST285 supporting an anti-inflammatory role for ST285. Upregulation of IFNγ, IL-1β and TNFα by monocytes following ST285 co-culture suggests a powerful defense against invading pathogens induced by ST285 that could be advantageous in defense against virus infection and tumours. Of interest, in spite of the upregulation of IFNγ, IL-1β and TNFα, considering collective down regulation of IFNAR1, IFNGR1, IL-18, our results might reveal an antagonistic effect of ST285 on pro-inflammatory IFNγ, IL-1β and TNFα responses which may lead to an overall downstream tolerance, and even an ultimate anti-inflammatory outcome.

### 4.3. ST285 activates mRNA expression of CXCL8 and downregulates CCR5 and CCL2

IL-8, also known as CXCL8 is produced by MQs; an important innate immune system chemokine which is associated with recruting neutrophils and other granulocytes of innate immune defense [50]. Our findings show a significant increased IL-8 gene expression by monocytes after exposure to ST285. We previously noted that ST1342, ST1275 and ST285 stimulate the U937 monocyte cell line to secrete increased levels of IL-8 [1]. Similarly, we showed PBMC exposure to ST285 results in overexpression of IL-8 [26]. Correspondingly, EPS from *L. paracasei* DG probiotic displayed immune-stimulating effects to human monocytic cell line THP-1 by increased expression of IL-8 gene [41]. In contrast, it was shown that dairy and soy fermented milks inoculated with *S. thermophilus* ST5 (ST5) mixed with either *L. helveticus* R0052 (R0052) or *B. longum* R0175 (R0175) added to LPS-challenged THP-1 monocyte cell line, decreased IL-8 production only when co-cultured with ST5+R0175 [51]. In addition, milk fermented with ST5+R0052 or ST5+R0175 did not alter the production of IL-8 by U937 monocyte cell line, whilst soy ferment prepared with ST5+R0175 downregulated IL-8 production [51].

C-C chemokine receptor type 5 (CCR5, CD195) and chemokine (C-C motif) ligand (CCL) 2 are mainly expressed on monocytes, DCs and MQs [52]. CCR5 is associated with Th1 immune responses and CCL2 with pathogenicity of a number of inflammatory diseases including rheumatoid arthritis and psoriasis, categorized by monocytic infiltrates through chemo-attracting monocytes [53]. Monocytes co-cultured with ST285 significantly downregulated CCR5 and CCL2, which is similar to ST285 co-cultured with PBMCs [26], suggesting an anti-inflammatory influence of ST285.

Although overexpression of IL-8 exclusively, may be interpreted as an inflammatory effect, taking into account the largely upregulated anti-inflammatory cocktail of cytokines and mediators induced by ST285, can in fact modulate this effect towards an anti-inflammatory profile for ST285. Upregulated IL-8 might be an initiating function of ST285 in order to trigger immune responses in the innate immune system, which then gets controlled by ST285 through reduction in the expression of CCR5. This in turn may lead to reduced Th1 immune responses, as well as decreased CCL2 and subsequently resulting in a controlled recruitment of monocyte. These effects may again highlight immunomodulatory effects of ST285 bacteria.

### 4.4. ST285 significantly upregulates mRNA expression level of colony stimulating factor

Colony stimulating factor (CSF, GM-CSF) is secreted by monocyte/MQs and supports and induces propagation, differentiation and production of different immune cells, mainly monocyte/MQs which are fundamental in responses againts infections. CSF is significantly increased (63.82 fold) by monocyte cultures in the presence of ST285, which is in alignment to our recent data showing increased CSF gene expression by PBMC co-cultured with ST285 [26]. The secretion of GM-CSF showed insignificant difference amongst immunized mouse spleen cells co-cultured with ST285 plus recall MBP_83–99_ peptide analog, compared to culturing cells with media alone or media plus recall MBP_83–99_ peptide analog [36]. In addition, ST1275, ST1342 and ST285 were also noted to induce high levels of GM-CSF production by U937 monocyte cell line [1]. It is known that G-CSF induces the development of IL-10-producing cells [54], hence, suggesting that ST285 may have an anti-inflammatory effect on the immune system.

### 4.5. ST285 downregulates mRNA expression levels of toll-like receptors

Toll-like receptors (TLRs) are mediators of innate immune responses primarily required in the defense against pathogens [55]. ST285 induced significant downregulation of TLR-1, TLR-2, TLR-4, TLR-5, TLR-6 and TLR-8, similar to our previous findings showing reduction of several TLRs by PBMC co-cultured with ST285[26]. Activated TLR (especially TLR-2 and TLR-4) together with other immune system factors can facilitate pro-inflammatory responses as well as further stimulating innate immune system actions [56-58]. Thus, an increased expression of TLR-2 and TLR-4 can lead to predominant inflammatory responses in the host, and their downregulation suggests reduction in such pro-inflammatory responses. Moreover, TLR-5 activation leads to stimulation of NF-κB which results in pro-inflammatory TNF-α production [59] and its reduced expression in monocyte co-cultured with ST285 may additionally signify an anti-inflammatory role for ST bacteria. Similar to our findings, another study has shown decreased expression of TLR-2, TLR-4, and TLR-9 using *L. plantarum* genomic DNA with THP-1 monocyte cells [48]. However, a study using human monocytes and monocyte-derived DCs exposed to UV-radiated *V. parvula, E. coli, B. adolescentis* and *L. plantarum*, showed higher expression of TLR-2 on monocytes compared to DCs, while TLR-4 was not detectable on DCs [42]. Additionally, in the same study it was shown that TLR-4 expression on monocytes was also down regulated in response to exposure to either *E. coli* or *L. plantarum* [42]. Downregulation in mRNA expression of TLRs genes, specifically when it occurs across a wide range including TLR-1, TLR-2, TLR-4, TLR-5, TLR-6 and TLR-8, designates anti-inflammatory properties for ST285.

### 4.6. ST285 downregulates cell surface markers CD14, CD86, CD4

CD14 is expressed on the cell surface of monocytes, MQs and DC and primarily binds to bacterial components [60-62], CD14 was significantly downregulated when co-cultured with ST285 bacteria suggesting an anti-inflammatory response. CD14 together with TLR-4 bind to bacterial components and both CD14 and TLR-4 were downregulated in the presence of ST285 bacteria. Co-culture of PBMC with ST285 also led to downregulated CD14 and TLR-4 expression [26]. However, ST285 upregulated expression of CD14 by U937 monocyte cell line [1]. In accord to our findings, human monocytes isolated from PBMC and exposed to *E. coli* or *L. plantarum* displayed down-regulated expression of CD14 [42]. CD86 (B7-2), is a co-stimulatory molecule necessary for initiating and maintaining T cells. Expression of CD86 mRNA levels by monocyte is significantly downregulated following culture with ST285, in line with our previous findings where CD86 was downregulated by bulk PBMC cultures [26]. Therefore, ST285 seem to induce an anti-inflammatory profile. Likewise, ST5+R0052 or ST5+R0175 milk or soy ferments also reduced expression of CD86 [51]. However, *L. fermentum* GR1485 and *L. plantarum* WCFS1 increased expression of CD86 by monocytes, inversely to *L. delbruekii* and *L. rhamnosus* that reduced CD86 expression [63]. Additionally, monocyte derived immature DCs co-cultured with *L. lactis* subsp. cremoris ARH74, *B. breve* Bb99 and *S. thermophilus* THS increased the expression of CD86 [23]. The contrast in these findings is not surprising and may be due to the dissimilarities in the nature of experiments; co-culture of monocytes with ST285 bacteria only compared to differentiated monocytes into immature DCs co-cultured with several probiotics or associated with differences in the properties of each bacteria.

CD4 is an extracellular cell surface molecule expressed by monocytes, MQ, DCs and Th cells and acts as a co-receptor between T cells and antigen presenting cells [64]. CD4 was significantly downregulated in monocyte cultures with ST285. In HIV-infected monocytes and MQs, CD4 is required for entry into the cell, and suggest that ST285 may have anti-viral properties.

Given the functional role of cell surface markers in immune responses, CD14 involvement in native immune responses, CD86 in T cell activation, and presence of CD4 on many cells underpinning innate and adaptive immunity, their downregulation in the presence of ST285 indicates an anti-inflammatory and anti-stimulatory profile for ST285. Additionally, due to the role of these cell surface markers in mediating innate and/or adaptive immune responses in defence against bacteria, downregulation of such markers could be suggestive of ST285 initiating self-tolerance via its immune modulation effects.

### 4.7. ST285 differentially downregulates mRNA expression level of other innate and adaptive immune response markers and chemokines

Integrin alpha M (ITGAM) or CD11b is another innate immune response factor associated with several inflammatory reactions such as phagocytosis, cell-mediated cytotoxicity, and chemotaxis. Lysozyme (LYZ) is also an innate immune response mediator associated with several inflammatory actions exists in mononuclear phagocytes such as MQs and performs as an antimicrobial enzyme. ITGAM and LYZ gene expressions are vastly downregulated in monocytes co-cultured with ST285, similarly to our recent findings showing downregulation of ITGAM and LYZ by PBMC co-cultured with ST285 [26]. Conversely, in U937 monocyte cell line, exposure to ST285 caused significant upregulation of CD11b/ ITGAM [1], the contrast could be related to difference between monocytes from healthy blood donors compared to monocyte cell line. MYD88, implicated in innate immunity, is downregulated by monocytes in response to ST285 co-culture. IL1RA1 has been shown to interact with MYD88 (together with PIK3R1 and IL1RAP) [65], is also downregulated, which both additionally highlight an anti-inflammatory role for ST285. Non-receptor tyrosine-protein kinase (TYK2) is an enzyme involved in various cellular events and extensive studies of TYK2-deficient mice indicate compromised IFNα, IL-12, and IL-23 pathways [66], and IL-12/Th1 and IL-23/Th17 axes [67], but it is dispensable for the signaling pathways of IL-6 or IL-10 [66]. It is believed that TKY2 is associated with a broader cellular pathways in human and it has a role in IL-12/Th1 and IL-23/Th17 axes involved in inflammatory/ autoimmunity, highlighting TKY2 choice as an effective therapeutic approach for select autoimmune diseases [66]. TYK2 is significantly downregulated by monocytes upon co-culture with ST285, a similar trend was found in our results when PBMC co-cultured with ST285 recently [26], which mutually support an anti-inflammatory profile for ST285.

IL-1-receptor-associated kinase-1 (IRAK1) is involved in innate immunity, and ST285 induced a significant downregulation of IRAK1 by monocyte culture. Likewise, *L. paracasei* stimulated the expression of IRAK3, but not IRAK1 in THP-1 cell line post differentiation with PMA. IRAK4 inhibitor suppressed the expression of negative regulators [40]. In contrast, THP-1 monocyte cells treated with *L. plantarum* genomic DNA induced a slight increase in IRAK-1 production [48]. SLC11A1 is a monocyte-MQ protein-1 involved in T cell activation and inflammatory disorders such as type 1 diabetes [68, 69], Crohn’s disease [70] and rheumatoid arthritis [71], is downregulated by monocytes upon co-culture with ST285. Our previous findings using ST285 to co-culture with PBMC similarly showed a reduced expression of SLC11A1[26], again suggesting an anti-inflammatory role for ST285. Induced downregulation of IRAK1, MYD88, TKY2, ITGAM, NOD2, SLC11A1 and LYZ by monocytes due to exposure to ST285 is suggestive of anti-inflammatory effects of ST285.

## 5. Conclusion

Commensal bacteria and probiotics have made their entry to the mainstream of healthcare and contribute to immune homeostasis in the gastrointestinal tract as well as conferring beneficial immunomodulatory properties that assist in the maintenance of a healthy immune system. ST is commonly applied in dairy products to ferment cheeses and yogurts and is thought to be beneficial to human health. We assessed the immune modulatory effects of ST285 on human monocytes and demonstrated that it delivers a range of potential immunomodulatory and anti-inflammatory properties. ST285 decreases mRNA expression of IL-1R, IL-18, IFNγR1, IFNαR1, CCL2, CCR5, TLR-1, TLR-2, TLR-4, TLR-5, TLR-6, TLR-8, CD14, CD86, CD4, ITGAM, LYZ, TYK2, IFNR1, IRAK-1, NOD2, MYD88, ITGAM, SLC11A1, and upregulates IL-1α, IL-1β, IL-2, IL-6, IL-8, IL-23, IFNγ, TNFα, CSF-2. No changes to mRNA expression were noted with IL-4, IL-5, IL-13, CCL2, CCL5, CCL8, CCR4, CCR8, CXCR3, CXCL10, TLR-3, TLR-9, CD8A, CD40, CD80, IFNB1, MPO, FOXP3, GATA3, STAT3, CD40LG, HLA-A, HLA-E, RORC. The data exhibits a predominant anti-inflammatory profile of cytokine, chemokine and cell markers induced by ST285.Therefore, the use of ST285 may be an efficacious approach for the treatment of select autoimmune diseases without using broad immunosuppression caused by currently available treatments for autoimmune disorders. Supplementary work is required to determine whether ST bacteria displays similar anti-inflammatory effects *in vitro* and *in vivo* in compromised immune disorders/ models such as inflammatory bowel disease, multiple sclerosis and allergies.

## Acknowledgements

All authors were supported by the Institute for Health and Sport and the Institute for Sustainable Industries and Liveable Cities at Victoria University. JJ was supported by a VU Research Fellowship, Victoria University Australia.

